# *Haloargentinum marplatensis* gen. nov., sp. nov., a novel extremely halophilic bacterium isolated from salted-ripened anchovy (*Engraulis anchoita*)

**DOI:** 10.1101/2020.07.27.213165

**Authors:** Silvina Perez, Margarita Gomila, Silvia Elena Murialdo, Irene Mabel Ameztoy, Narjol Gonzalez-Escalona, Elida Elvia Ramírez, María Isabel Yeannes

## Abstract

A facultative aerobic, Gram-negative, motile, non-endospore forming and extremely halophilic bacterium, strain 11aii^⊺^, isolated from salted-ripened anchovy, was examined using a polyphasic approach to characterize and clarify its phylogenetic and taxonomic position. Sequences of the 16S rRNA gene revealed close relationships to species of the genera *Lentibacillus* and *Virgibacillus* (94.2% similarity). The organism grew optimally in the presence of 20-35 % NaCl. The major fatty acids of strain 11aii^⊺^ were C_16:0_ (42.1%) and anteiso-C15_:0_ (31.2%) and also presented iso-C_16:0_ (11.0%), anteiso-C_17:0_ (10.4%) and C_18:0_ (5.2%). Based on data presented here, strain 11aii^⊺^ is considered to represent a novel genus and species, for which the name *Haloargentinum marplatensis gen*. nov. sp. nov. is proposed with the strain 11aii^⊺^ as type strain.

## Introduction

Salting is an ancient method that has been applied for fish preservation. It can be followed by the ripening stage consisting of chemical and physicochemical changes that modify the characteristics of the muscle tissue and thus the sensory properties of the fish. These changes require months in the presence of high salt content (NaCl). Salting and ripening of different pelagic species is a worldwide common and traditional practice [1, 2]. Among this type of product, salted-ripened anchovy (*Engraulis anchoita*) produced in Latin American countries can be mentioned. Due to the high NaCl content and low water activity values that characterize this type of products, the microbiota is mainly constituted by halophilic or halotolerant microbes. The role of microorganisms during the ripening is under continuous investigation [1, 3–7]. Recent studies have reported the isolation of many novel bacteria and archaea from salted and fermented seafood: *Lentibacillus jeotgali* Grbi^⊺^ [8], *Halomonas shantousis* SWA25^⊺^ [9], *Halobacterium piscisalsi* HPC1-2^⊺^ [10], *Natrinema gari* HIS40-3^⊺^ [11], *Haloterrigena jeotgali* A29^⊺^ [12], *Haloarcula salaria* HST01-2R^⊺^ and *Haloarcula tradensis* HST03^⊺^ [13]. Here, we report the taxonomic characterization of an halophilic isolate which closest relatives are members of the genera *Lentibacillus* and *Virgibacillus* belonged to the family *Bacillaceae* [8,14, 15]. The genus *Lentibacillus* was defined by Yoon et al. [16], with the description of *Lentibacillus salicampi* SF-20^⊺^, a Gram-variable endospore-forming rod-shaped strain. Its last described species corresponds to *Lentibacillus lipolyticus* SSKP1-9^⊺^ isolated from salted shrimp paste in Thailand [17]. The genus *Virgibacillus* was established by the reclassification of *Bacillus pantothenticus* CN3028^⊺^ [18] as *Virgibacillus pantothenticus* [19], and the genus description was later emended by Heyrman et al [20]. Members of this genus are Gram-positive or Gram-variable, endospore-forming, motile rods [20, 21]. At the time of writing, *Lentibacillus* and *Virgibacillus* genera contained 17 and 36 validly named species, respectively, as reported on the LSPN website (www.bacterio.net/lentibacillus.html and www.bacterio.net/virgibacillus.html). Notably, the reported halophilic isolate here presented remarkable morphotype differences with the mentioned genera. Based on the results of our taxonomic study and previous characterizations of the most closely related genera, we consider that the halophilic strain should be included within a novel genus and species for which the name *Haloargentinum marplatensis* gen. nov., sp. nov. is proposed.

## Isolation and Ecology

Strain 11aii^⊺^ was isolated from beheaded and partially gutted salted-ripened anchovies collected from a local factory (Mar del Plata, Argentina). Homogenates were prepared in saline broth (NaCl, 150 g/L; meat peptone, 3 g/L; meat extract, 3 g/L) in duplicate, followed by a subsequently enrichment step performed by incubation at 35–37 °C for 60 min and successive serial dilutions were carried out [22]. Homogenates (0.1 mL) were spread onto the growth media named Tryptone-salt-yeast extract (TSL: NaCl, 200 g/L; MgSO_4_(7H_2_O), 20 g/L; KCl, 5 g/L; CaCl_2_(6H_2_O), 0.2 g/L; tryptone, 5 g/L; yeast extract, 4 g/L; agar-agar, 17 g/L) [23] in duplicate and incubated at 35–37 °C during 21 days. Colonies with different macroscopic characteristics (colour, size, shape and density) were re-streaked on fresh agar plates and incubated at 35-37 °C until growth. Pure isolates were transferred to TSL broth (NaCl, 200 g/L; MgSO_4_(7H_2_O), 20 g/L; KCl, 5 g/L; CaCl_2_(6H_2_O), 0.2 g/L; tryptone, 5 g/L; yeast extract, 4 g/L) [23]. Colony stocks were kept at 4 °C for further analyses.

## 16S RNA phylogeny

DNA was extracted and purified as described by D’Ippólito *et al.* [24] and Sheu *et al.* [25]. The reaction mixture for PCR was performed with 5 μL of DNA template (cell-by-heat lysate), 2.5 μL of buffer 1X, 1.5 μL of MgCl2 50 mM, 1.25 μL of dimethyl-sulphoxide, (DMSO), 1.25 μL dNTPs 10 mM, 0.8 μL of each primer (F43Eco 5’-CGGAATTCCAGGCCTAACACATGCAAGTC-3’ and R1387Eco 5’-CGGAATTCGGGCGGWGTGTACAAGGC-3’), 0.25 μL of Taq polimerase, to a final volume of 25 μL. PCR reaction was executed by Biometra UNO-Thermoblock Thermal Cycler. Amplifications were carried out using the following program: (94 °C 3 min) x 1; (94 °C 1 min, 55 °C 1 min, 72 °C 90 s) x 30, (72 °C 10 min) x 1. PCR products were purified by QIAquick PCR Purification kit (Qiagen, Alemania). PCR products (10 μl each) were analyzed on 2 % TAE pre-cast agarose gels (Bio-Rad, Hercules, CA) and run at 75 V for 1 h in 1X TAE with a molecular weight standard (100 bp ladder, Promega, WI, USA). Amplification products were visualized by ethidium bromide staining (5 ug/ml). PCR product consisted of a single band. PCR product was sequenced in both directions by MCLAB (South San Francisco, CA, USA) employing primers 27F, 357F (5’-CTCCTACGGGAGGCAGCAG-3’), 518R (5’-CGTATTACCGCGGCTGCTGG-3’), and 1492R sequenced by MCLAB company (www.mclab.com). DNA sequences were assembled using Bioedit [26].

For 16S rDNA phylogenetic analysis, a BLAST analysis of the 11aii^⊺^ strain 16S rDNA sequence showed that it matched 94 % to 16S rDNA sequences from strains *Lentibacillus* sp. KM1091 and *Lentibacillus juripiscarius* strain P1-ASH. Ninety eight 16S rDNA partial sequences representing 90 highly related bacterial taxa (publicly available at GenBank - Supplementary Table 1) and with a 92-99 % identical to the 16S rDNA from strain 11aii^⊺^ were retrieved in order to re-construct the phylogenic relationship of the strains. The sequences were aligned and an internal 16S rDNA fragment of 1190 bp was used for the phylogenetic study. The phylogeny was reconstructed using the maximum likelihood method using Mega software v7 [27] and using the kimura-2 parameter model to estimate the genetic distances [28]. The statistical support of the nodes in the ML tree was assessed by 500 bootstrap re-sampling.

Figure 1 shows the phylogenetic tree with the most of the Lineages compressed for visualization purposes. The original phylogenetic tree with the highest log likelihood is shown in Supplementary Figure 1. The 16S rRNA gene sequence of strain 11aii^⊺^ was closely related to species from the genera *Lentibacillus* and *Virgibacillus* (94.2% similarity) by phylogentic analysis. It was closely related to *Lentibacillus juripiscarius* (93.4%), *Lentibacillus jeotgali* (92.2%), *Virgibacillus flavescens* (92.0%) and *Virgibacillus phasianinus* (91.7%).

**Figure 1.**
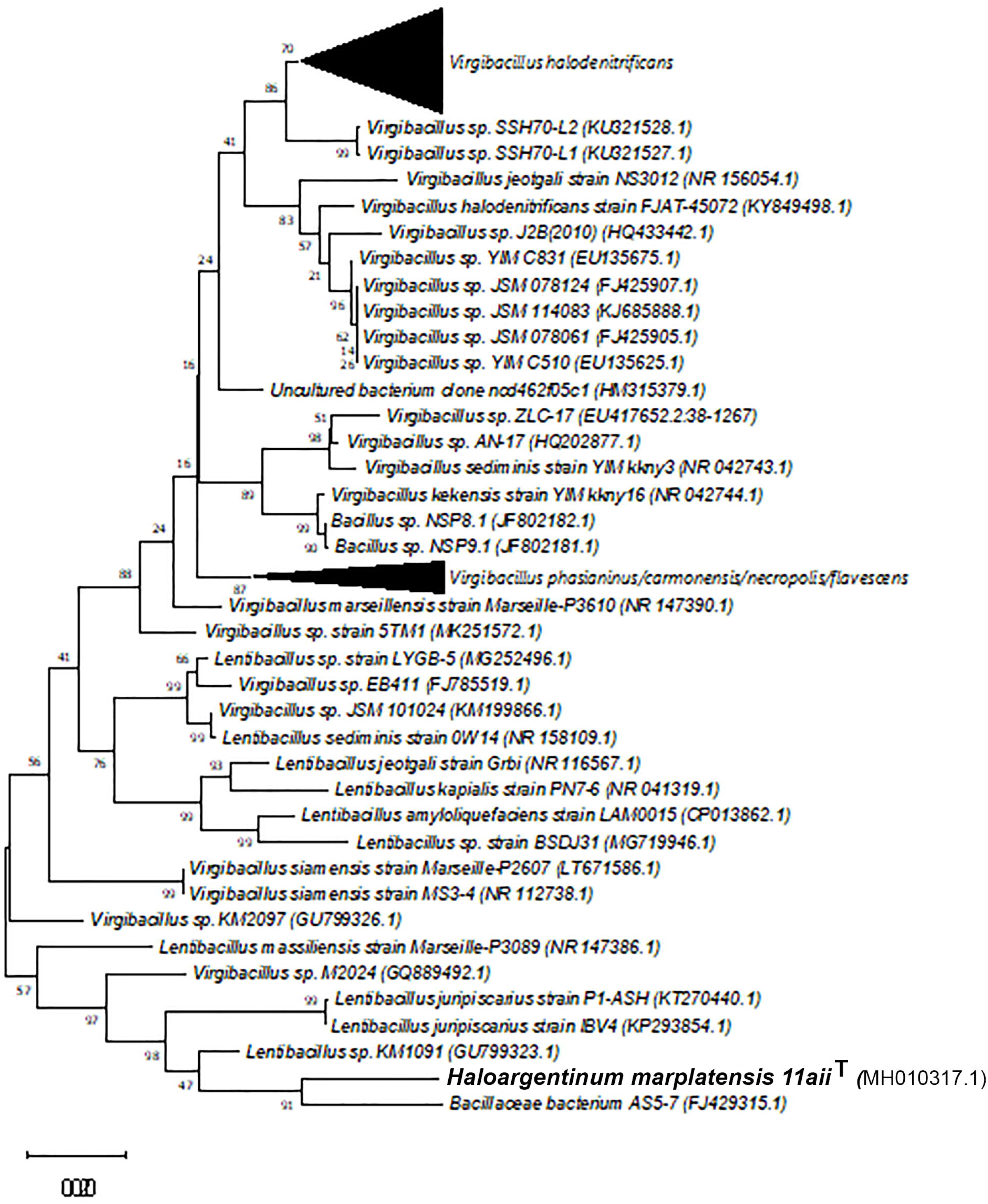
Maximum likelihood phylogeny of strain 11aii^⊺^ with closest relatives using a 1190 bp fragment of the 16S rDNA gene. The evolutionary history was inferred by using the Maximum Likelihood method based on the Kimura 2-parameter model [28]. The tree shown has most of the Lineages compressed for visualization purposes. Bootstrap support above 50% are shown above the branches. In red fonts are the strains sequenced in this study. The tree is drawn to scale, with branch lengths measured in the number of substitutions per site. Evolutionary analyses were conducted in MEGA7 [27].

## Physiology and Chemotaxonomy

Morphological, physiological and biochemical characteristics were studied. The cell morphology was carried out on the basis of the Gram staining (optic microscope) [29]. Focused on the capacity to produce histamine by decarboxylation of its precursor (histidine), the histidine-decarboxylase test was carried out. Cultures were inoculated on slanting surface of a solidified selective medium (tryptone, 5 g/L; yeast extract, 5 g/L; L-histidine, 27 g/L; CaCO_3_, 1 g/L; agar-agar, 20 g/L; bromocresol purple, 0.06 g/L; pH 5.3), and incubated at 35–37°C during 10 days. Positive result was indicated by the medium towards violet [30, 31]. Isolate with positive histidine-decarboxylase test was submitted to a further characterization. Therefore, Ziehl-Neelsen staining was carried out and spore staining was done by the Schaeffer and Fulton [32] technique. NaCl requirement was determined in the above growth broth containing various NaCl concentrations (0–6 M). Inoculums were incubated at 35–37 °C and positive result was indicated by growing. Growth at different pH values (5.0 to 8.5, with an interval of 0.5) was examined using TSL. Tests for catalase and cytochrome oxidase activities, motility, nitrate reduction, urease reaction, lysine decarboxylase in lysine iron agar, histidine and arginine dihydrolase by inoculation in basal broth with the respective amino acid, citrate utilization on Simmons citrate agar (Britania) and the hydrolysis of gelatine and starch were performed as described by MacFaddin [29]. The hydrolysis of Tween 80 was detected by screening for zones of hydrolysis around colonies growing in a solid medium containing 1% v/v of this subtract [33]. Hydrogen sulphide (H_2_S) production was tested by inoculation in TSI medium (Britania) which allows the investigation of the production of H_2_S and also the production of acid and gas from glucose, lactose and sucrose. Indole formation was studied by inoculation and growing in peptone broth and subsequently reaction with Kovacs’s reagent. Acid production from carbohydrates was determined in red phenol broths with 1% w/v of each subtract under study (galactose, sucrose, glucose, fructose, lactose, maltose, sorbitol, mannitol, trehalose, xylose and arabinose). Oxidative/fermentative metabolism of glucose was determined on OF basal medium [29]. Proteolytic and lipolytic activities were determined by streaking pure culture in skim milk agar (yeast extract, 3 g/L; meat peptone, 5 g/L; agar, 15 g/L; milk, 10 mL/L) and in a solid medium containing 1% v/v of tributyrin, respectively. Inoculated plates were incubated at 35–37 °C for 10 days. Clear zones around the streaks were regarded as positive reactions [34]. All culture media used for biochemical tests were supplemented with NaCl to a final concentration of 200 g/L, with K^+^ (10 ppm) and Mg^2+^ (0.1 ppm) in order to provide the specific nutrients needed by halophilic bacteria [23]. All analyses were carried out in duplicate.

Based on colony macroscopic characteristics, two isolates were distinguished, namely, 11ai and 11aii^⊺^ llai colonies were pale-pink pigmented, their cells were Gram-negative long-rods-shaped and this isolate resulted negative for histidine-decarboxylase test. The 11aii^⊺^ colonies formed on agar plates were circular (1–2 mm in diameter), smooth, translucent and salmon-reddish pigmented. This isolate was Gram-negative and cells coccobacilli and disc-shaped (pleomorphic) were observed. Regarding to the histidine-decarboxylase test, 11aii^⊺^ was positive, indicating that it could form histamine. The presence of this biogenic amine is regulated because of in high concentrations represents a potential food safety hazard [35, 36]. Therefore, 11aii^⊺^ was selected for further investigations. Ziehl-Neelsen staining exhibited a negative result. Endospores was not observed. The strain was able to grow at high NaCl concentrations, from 3.4 M (approximately 20 %) to 6 M (approximately 35 %), and pH between 5.5 and 8. The strain was positive for oxidase and catalase. Acid was not produced from sugars by red phenol broths, TSI or OF basal medium. Cells did not hydrolyse gelatine or Tween 80 but they did starch. The strain was positive for nitrate reduction and indole formation in the presence of tryptophan and but negative for urease reaction, citrate utilization and hydrogen sulphide production. This isolate was positive for histidine decarboxylase but negative for lysine decarboxylase and arginine dihydrolase. The strain was lipolytic and non-proteolytic by the method of FIL IDF 73 [34], i.e. tributyrin was hydrolysed but casein was not. Characteristics that distinguish isolate 11aii^⊺^ from recognized members of the genus *Lentibacillus* and *Virgibacillus* are summarized in Table 1. The new strain can be differentiated from other closely related species by several phenotypic properties, noting that it is Gram-negative, pleomorphic, absent endospores and no sugars fermenter.

**Table 1.** Differential phenotypic characteristics between strain *Haloargentinum marplatensis gen*. nov. sp. nov. 11aii^⊺^ and species of the closely related *Lentibacillus* and *Virgibacillus* genera. Strains: 1, 11aii^⊺^ (data from the present study); 2, *Lentibacillus jeotgali* Grbi [8]; 3, *Lentibacillus juripiscarius* IS40-3^⊺^ [39]; 4, *Lentibacillus massiliensis* Marseille-P3089^⊺^ [40]; 5, *Virgibacillus flavescens* S1-20^⊺^ [41]; 6, *Virgibacillus halodenitrificans* [14]; 7, *Virgibacillus kekensis YIM* kkny16^⊺^ [42]; 8, *Virgibacillus phasianinus* LM2416^⊺^ [15]; 9, *Virgibacillus siamensis* MS3-4^⊺^ [43]. Symbols: +, positive reaction; –, negative reaction; w, weakly positive; v, variable; ND, no data.

| Characteristic | 1 | 2 | 3 | 4 | 5 | 6 | 7 | 8 | 9 |
| --- | --- | --- | --- | --- | --- | --- | --- | --- | --- |
| Isolation source | Salted-ripened anchovies, Argentina | Fermented scallops, Korea | Fish sauce, Thailand | Salty stool, Senegal | Marine sediment, China | Marine solar saltern, Korea | Salt lake, China | Faeces of *Lophura swinhoii*, Korea | Fermented fish, Thailand |
| Pigmentation | Salmon-reddish | - | - | yellow | Light yellow | - | creamy grey | - | red color |
| Gram staining | - | + | + | + | v | V | + | + | + |
| Cell morphology | Pleomorphic | Rods | Rods | ND | Rods | Rods | Rods | Rods | Rods |
| Endospore forming | - | + | + | + | ND | + | + | ND | + |
| Motility | + | - | - | ND | + | + | + | + | + |
| NaCl range (% w/v) | 20-35 | 3-20 | 3-30 | 0.5-20 | 0-20 | 2-25 | 0-25 | 0-20 | 1-20 |
| pH range | 5.5-8.0 | 6.0-8.0 | 5.0-9.0 | ND | 7.0-9.0 | 5.8-9.6 | 6.0-10.0 | 6.0-7.0 | 5.0-8.0 |
| Growth at 35 °C | + | + | + | ND | - | + | + | - | + |
| Nitrate reduction | + | + | + | ND | - | + | + | + | - |
| Indole formation | + | ND | - | ND | - | - | - | ND | ND |
| Oxidase | + | - | + | + | + | + | + | - | + |
| Catalase | + | + | + | - | + | + | + | + | + |
| Urease reaction | - | - | - | ND | - | - | - | - | ND |
| Lysine decarboxylase | - | ND | ND | ND | - | - | ND | + | ND |
| Arginine dihydrolase | - | - | - | ND | - | - | ND | - | ND |
| Citrate utilization | - | ND | ND | ND | - | ND | ND | + | - |
| H<sub>2</sub>S production | - | ND | - | ND | - | - | - | ND | - |
| Acid production from: |  |  |  |  |  |  |  |  |  |
| Galactose | - | + | + | - | ND | + | - | + | - |
| Sucrose | - | - | + | W | ND | + | - | - | - |
| Glucose | - | + | + | + | ND | + | + | + | - |
| Fructose | - | + | + | + | ND | + | - | + | - |
| Lactose | - | - | v | - | ND | + | - | + | - |
| Maltose | - | + | + | - | ND | + | + | + | - |
| Sorbitol | - | ND | - | - | ND | + | - | ND | ND |
| Mannitol | - | + | v | - | ND | + | w | ND | - |
| Trehalose | - | - | + | - | ND | + | w | - | - |
| Xylose | - | - | - | + | ND | - | - | - | - |
| Arabinose | - | - | - | - | ND | - | - | - | - |
| Hydrolysis of |  |  |  |  |  |  |  |  |  |
| Gelatin | - | ND | + | + | ND | ND | - | + | + |
| Starch | + | ND | - | - | ND | ND | + | + | - |
| Tween 80 | - | - | - | + | ND | ND | - | - | - |
| Tributyrin | + | ND | ND | - | ND | ND | ND | ND | ND |
| Casein | - | - | + | + | ND | ND | - | + | + |

For cellular fatty acid analysis, strain was cultured on halophilic growth broth for a week at 35 °C and the fatty acids were extracted as described by Bligh and Dyer [37] and the extract was dried under nitrogen gas. The determination of Fatty acid methyl esters (FAME) profile was realized by gas chromatography coupled to mass spectrometry (GC-MS) using 2% sulphuric acid–methanol (v/v) as methylating reagent and methyl-nonadecanoate as internal standard [38]. The Thermo Scientific TRACE 1300 Mainframe MS 230V gas chromatograph was used with the TG-5MS column (0.25 mm, 30 m, Thermo Scientific) coupled to the Thermo Scientific ISO. mass detector (single quadrupole) with vacuum closing system. The GC-MS program consisted of programmed temperature vaporizer (PTV) at 200°C, flow rate of 40.5 mL/min with split ratio 1/45 and oven temperature of 160 °C maintained for 5 min, 5°C/min up to 300°C and maintained 5 min. The relative amount of each CFA was expressed as percentage of the total fatty acids. The fatty acids of strain 11aii^⊺^ were C_16:0_ (42.1%) and anteiso-C_15:0_(31.2%) and also presented iso-C_16:0_ (11.0%), anteiso-C_17:0_ (10.4%) and C_18:0_ (5.2%). Comparison of CFA profile of the strain 11aii^⊺^ and closely related is indicated in Table 2. As in other species of related genera, Anteiso-C_15:0_, Iso-C_16:0_ and Anteiso-C_17:0_ represented an important proportion of the cellular fatty acids. However, 11aii^⊺^ major fatty acid was C_16:0_ differentiating from the other species where it did not exceed 3%.

**Table 2.** Comparison of fatty acid compositions between characteristics between strain *Haloargentinum marplatensis* gen. nov. sp. nov. 11aii^⊺^ and species of the closely related *Lentibacillus* and *Virgibacillus* genera. Strains: 1, 11aii^⊺^ (data from the present study); 2, *Lentibacillus jeotgali* Grbi^⊺^ [8]; 3, *Lentibacillus juripiscarius* IS40-3^⊺^ [39]; 4, *Virgibacillus flavescens* S1-20^⊺^ [41]; 5, *Virgibacillus halodenitrificans* [14]; 6, *Virgibacillus kekensis YIM* kkny16^⊺^ [42]; 7, *Virgibacillus phasianinus* LM2416^⊺^ [15]; 8, *Virgibacillus siamensis* MS3-4^⊺^ [43];. Data are percentages of the total fatty acids; components representing less than 1.0% of the total are not shown.

| Fatty acid | 1 | 2 | 3 | 4 | 5 | 6 | 7 | 8 |
| --- | --- | --- | --- | --- | --- | --- | --- | --- |
| *Saturated* |  |  |  |  |  |  |  |  |
| *C*<sub>14:0</sub> |  |  |  |  | 1.7 |  |  |  |
| *C*<sub>16:0</sub> | 42.1 |  |  | 1.2 | 2.7 |  |  | 1.5 |
| *C*<sub>18:0</sub> | 5.2 |  |  |  | 2.1 |  |  |  |
| *Unsaturated* |  |  |  |  |  |  |  |  |
| *C*<sub>16:1</sub>ω7c alcohol |  |  | 2.6 |  | 8.1 |  |  |  |
| *C*<sub>18:1</sub>ω9c |  |  |  |  | 1.1 |  |  |  |
| *Branched* |  |  |  |  |  |  |  |  |
| *Iso-C*<sub>14:0</sub> |  | 5–13 | 1 | 18.2 | 13.3 |  | 3.1 | 3.9 |
| *Iso-C*<sub>15:0</sub> |  | 3–18 | 4.4 | 2.1 | 2.6 |  | 5.3 | 11.3 |
| *Anteiso-C*<sub>15:0</sub> | 31.2 | 38–54 | 61.9 | 30.3 | 50.4 | 54.1 | 73.0 | 55.8 |
| *Iso-C*<sub>16:0</sub> | 11.0 | 13–30 | 4.5 | 36.4 | 6.1 |  | 5.2 | 6.6 |
| *Iso-C*<sub>17:0</sub> |  |  |  |  |  |  |  | 1.5 |
| *Anteiso-C*<sub>17:0</sub> | 10.4 | 13–18 | 20 | 9.8 | 7.0 | 32.0 | 9.7 | 17.7 |
| Summed feature* |  |  |  |  |  |  |  |  |
| 4 |  |  | 2.5 |  |  |  |  |  |

\*Summed feature 4 comprises anteiso-C17:1 B and/or iso-C17:1 I

In conclusion, results of phenotypic, genotypic and phylogenetic studies presented in this study demonstrate that strain 11aii^⊺^ represents a novel genus and species for which the name *Haloargentinum marplatensis* gen. nov., sp. nov. is proposed as a new representative of the phylum *Firmicutes*. Strain 11aii^⊺^ is the type strain of *Haloargentinum marplatensis*.

## Protologue

### Description of *Haloargentinum* gen. nov

*Haloargentinum* (Ha.lo.ar.gen.ti.num. Gr. masc. n. hals, *halos*, salt; N.L. neut. adj. argentina, pertaining to Argentina, where the bacteria was isolated; N.L. neut. n. *Haloargentinum*, salt (-requiring) and Argentina). Cells are Gram-negative, coccobacilli/Disc-shaped or pleomorphic bacteria, phylogenetically affiliated in the phylum *Firmicutes*. Aerobic. Oxidase- and catalase-positive. Extremely halophilic, requiring at least 200 g salt / L for growth. Habitat: salted and ripened anchovies. The type species is *Haloargentinum marplatensis*.

### Description of *Haloargentinum marplatensis* sp. nov

Cells are motile, Gram-negative coccobacilli/disc-shaped (pleomorphic) without endospores. Colonies formed on agar plates are circular (1–2 mm in diameter), smooth, translucent and salmon-reddish pigmented. Growth occurs in the presence of 20–35 % (w/v) NaCl and pH 5.5-8. The isolate is positive for oxidase and catalase and negative for Ziehl-Neelsen staining. Acid is not produced from carbohydrates (galactose, sucrose, glucose, fructose, lactose, maltose, sorbitol, mannitol, trehalose, xylose and arabinose). Cells hydrolyse starch but no gelatine and Tween 80. Positive for nitrate reduction and indole formation in the presence of tryptophan and negative for urease reaction, citrate utilization and hydrogen sulphide production. Histidine decarboxylase is present and lysine decarboxylase, arginine dihydrolase and phenylalanine deaminase are absent. Tributyrin hydrolysis is produced but no milk proteolysis (casein hydrolysis). Major fatty acids are n-C_16:0_ and anteiso-C_15:0_.

The type strain is 11aii^⊺^, was isolated from salted-ripened anchovies, a traditional fermented food elaborated in Argentina. The GenBank/EMBL/DDBJ accession number for the 16S rRNA gene sequence of strain 11aii^⊺^ is MH010317.

## AUTHOR STATEMENTS

### Authors and contributors

Silvina Perez: Investigation, Visualization, Writing-original draft. Margarita Gomila: Visualization, Writing – review & editing. Silvia Elena Murialdo: Funding acquisition, Supervision. Irene Mabel Ameztoy: Investigation. Narjol Gonzalez-Escalona: Software, Resources. Elida Elvia Ramírez: Investigation. María Isabel Yeannes: Conceptualization, Funding acquisition.

## Acknowledgements

Narjol Gonzalez Escalona was supported by the FDA Foods Program Intramural Funds. Silvina Perez, Irene Mabel Ameztoy and María Isabel Yeannes were supported by Consejo Nacional de Investigaciones Científicas y Técnicas (CONICET).

---

Supplementary figure 1. Phylogenetic tree with the highest log likelihood phylogeny of strain llaiiT with closest relatives using an 1190 bp fragment of the 16S rRNA gene. The evolutionary history was inferred by using the Maximum Likelihood method based on the Kimura 2-parameter model [28]. Bootstrap support above 50% are shown above the branches. In red fonts are the strains sequenced in this study. The tree is drawn to scale, with branch lengths measured in the number of substitutions per site. Evolutionary analyses were conducted in MEGA7 [27],

