## Supplementary figures and images for "*Haloargentinum marplatensis* gen. nov., sp. nov., a novel extremely halophilic bacterium isolated from salted-ripened anchovy (*Engraulis anchoita*)"

### Supplementary Figure 1

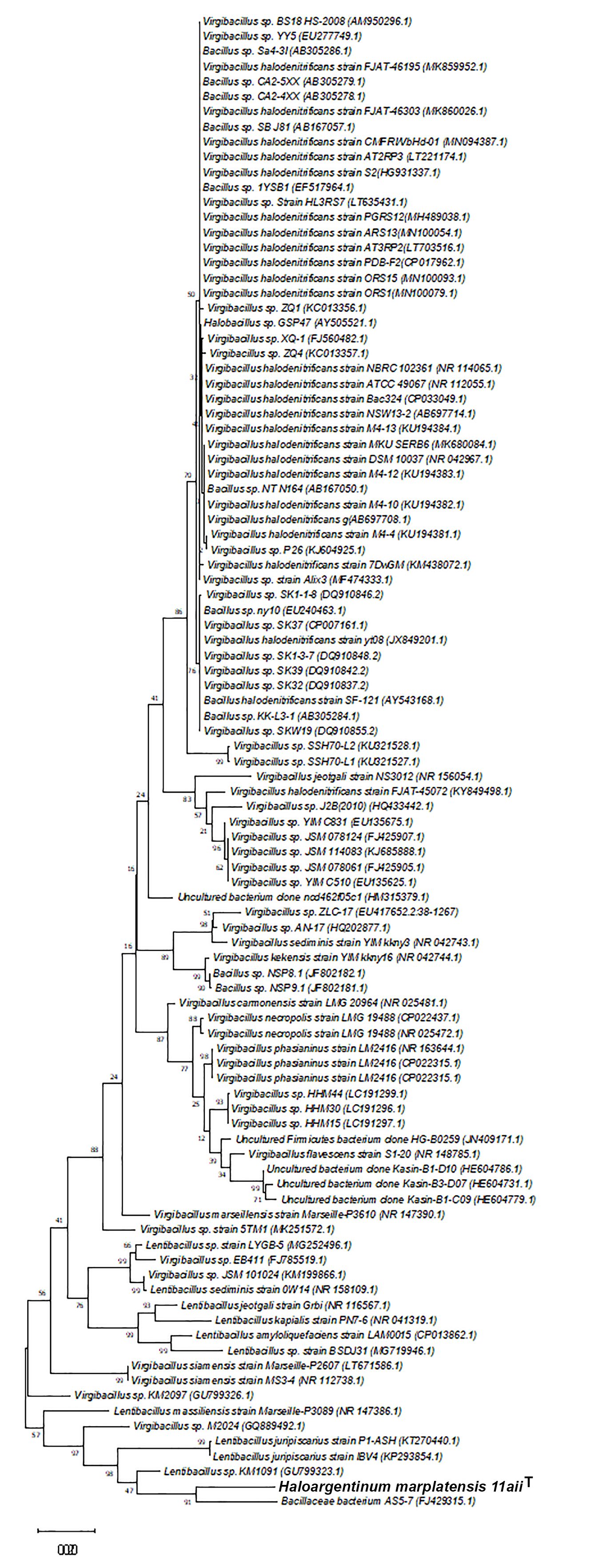
